# Rab11A regulates the constitutive secretory pathway during *Toxoplasma gondii* invasion of host cells and parasite replication

**DOI:** 10.1101/782391

**Authors:** Venugopal Kannan, Chehade Sylia, Werkmeister Elisabeth, Barois Nicolas, Periz Javier, Lafont Frank, Tardieux Isabelle, Khalife Jamal, Gordon Langsley, Meissner Markus, Marion Sabrina

**Affiliations:** Centre d’Infection et d’Immunité de Lille, Université de Lille, INSERM U1019, CNRS UMR 8204, Institut Pasteur de Lille, Lille, France; Institute for Advanced Biosciences (IAB), Membrane Dynamics of Parasite-Host Cell Interactions, CNRS UMR5309, INSERM U1209, Université Grenoble Alpes, Grenoble, France; Laboratoire de Biologie Cellulaire Comparative des Apicomplexes, Faculté de Médicine, Université Paris Descartes—Sorbonne Paris Cité, France. INSERM U1016, CNRS UMR8104, Institut Cochin, Paris, France; Department of Veterinary Sciences, Experimental Parasitology, Ludwig-Maximilians-Universität, Munich, Germany

**Author notes:** Wellcome Centre for Integrative Parasitology, Institute of Infection, Immunity and Inflammation, University of Glasgow, Glasgow, UK. contributed equally (Venugopal Kannan, Chehade Sylia).

## Abstract

*Toxoplasma gondii* possesses an armada of secreted virulent factors that enable parasite invasion and survival into host cells. These factors are contained in specific secretory organelles, the rhoptries, micronemes and dense granules that release their content upon host cell recognition. Dense granules are secreted in a constitutive manner during parasite replication and play a crucial role in modulating host metabolic and immune responses. While the molecular mechanisms triggering rhoptry and microneme release upon host cell adhesion have been well studied, constitutive secretion remains a poorly explored aspect of *T. gondii* vesicular trafficking. Here, we investigated the role of the small GTPase Rab11A, a known regulator of exocytosis in eukaryotic cells. Our data revealed an essential role of Rab11A in promoting the cytoskeleton driven transport of DG and the release of their content into the vacuolar space. Rab11A also regulates transmembrane protein trafficking and localization during parasite replication, indicating a broader role of Rab11A in cargo exocytosis at the plasma membrane. Moreover, we found that Rab11A also regulates extracellular parasite motility and adhesion to host cells. In line with these findings, MIC2 secretion was altered in Rab11A-defective parasites, which also exhibited severe morphological defects. Strikingly, by live imaging we observed a polarized accumulation of Rab11A-positive vesicles and dense granules at the apical pole of extracellular motile parasites suggesting that a Rab11A-dependent apically polarized transport of cargo regulates parasite motility.

## Introduction

*Toxoplasma gondii* (*T. gondii*) is an obligatory intracellular parasite that belongs to the phylum *Apicomplexa*, typified by the presence of specific apical secretory organelles, the rhoptries and the micronemes. Upon contact with the host cell, rhoptry (ROP) and microneme (MIC) proteins are released and promote parasite entry by driving the formation of a tight parasite-host cell adhesive membrane structure (called the circular junction) [1]. ROP proteins also contribute to building the parasitophorous vacuole (PV), within which the parasite rapidly replicates. The molecular mechanisms regulating MIC exocytosis have been well studied leading to the discovery of specific parasite signaling pathways triggering their secretion upon parasite adhesion to host cells [2] [3]. Dense granules (DG) are also parasite secretory organelles essential for parasite survival, which release effectors modulating host immune and metabolic responses [4]. Dense granule proteins (GRA) also promote the formation of an intravacuolar nanotubular network (IVN), which interconnects parasites during intracellular replication, thereby ensuring the synchronicity of the successive divisions [5] [6] [7]. The IVN also connects the parasite to the PV membrane (PVM), presumably enhancing parasite exchanges with its host, notably for nutrient retrieval and parasite effector release into the host cytosol [8]. In contrast to micronemes and rhoptries, DGs are randomly distributed in the parasite cytosol and the mechanisms regulating their exocytosis at the parasite plasma membrane (PM) have not been elucidated. In Metazoan, the process of exocytosis implies the active transport of secretory vesicles to the PM, and the secretion of their content into the extracellular environment or their insertion into the PM. In mammalian cells and plants, two different exocytic routes have been described: the “constitutive secretory pathway” supports the sorting of newly synthesized proteins from the endoplasmic reticulum, through the Golgi apparatus to the PM, while, the “recycling pathway” targets to the cell surface internalized material that has been transported to and sorted in the pericentriolar or peripheral recycling endosomes [9]. In *T. gondii*, DGs are considered to be the default constitutive secretory pathway based on the observation that the SAG1-GFP fusion protein (full product or truncated of its GPI anchor (SAG1ΔGPI)) was found to be transported by DGs before to being released into the vacuolar space [10]. In addition, proteins whose specific motifs targeting them to other secretory organelles have been deleted are localized in DGs [11] [12]. Yet, there is so far no evidence that transmembrane proteins navigate through the DGs to reach the PM of the parasite.

The small Rab GTPases belong to the Ras small G protein subfamily and operate as molecular switches that alternate between two conformational states: the GTP-bound “active” form and the GDP-bound “inactive” form [13]. Through their interactions with various effectors, such as coat components, molecular motors and soluble NSF attachment protein receptors (SNAREs), the Rab GTPases serve as multifaceted organizers of almost all membrane trafficking processes, including vesicle budding from the donor compartment, vesicle transport along cytoskeleton tracks and vesicle tethering and fusion at the acceptor membrane [13] [14] [15]. Among the Rab GTPases, Rab11 regulates the constitutive secretory and recycling pathways, thus controlling secretion at the PM [14] [15] [16]. In mammalian cells, Rab11 regulates vesicle transport via their anchoring to both, microtubule [17] and actin-based molecular motors [18]. In addition, Rab11A promotes the tethering of recycling vesicles to the PM in concert with the exocyst complex [19] [20] [21]. Through multiple interactions with effector molecules, Rab11 influences numerous cellular functions including ciliogenesis [22], cytokinesis [23] and cell migration [24] [25]. In contrast to humans, which express over 70 Rabs, *T. gondii* possesses a limited number of 13 Rabs that include two isoforms of Rab11, Rab11A and Rab11B [26]. In *T. gondii* as well as in the related apicomplexan geni *Plasmodium spp.*, Rab11A-defective parasites are unable to complete cytokinesis and show marked defects in the exocytosis-assisted process that leads to proper individualization of daughter cells, otherwise posteriorly connected [27] [28] [29]. In *Plasmodium*, this process was suggested to be regulated by a PI4K-Rab11A mediated secretion of vesicles from the TGN to the PM [28] [29]. Here, we further explored the functions of Rab11A in *T. gondii* and demonstrated its key role in the regulation of DG exocytosis and transmembrane protein delivery at the parasite PM. We also unraveled a novel role for Rab11A in extracellular parasite adhesion and motility, thereby contributing to host cell invasion.

## Results

### Rab11A localizes to dynamic cytoplasmic vesicles

To investigate *T. gondii* Rab11A localization, we raised a polyclonal antibody in mice, which recognized a unique protein at the expected size of 25kDa in a total extract of type I RHΔ*Ku80* parasites (Fig 1A). Next, we performed immunofluorescence assays (IFA) in fixed RHΔ*Ku80* tachyzoites. Rab11A displays distinct localizations depending on the cell cycle stage. During the G1 phase, Rab11A is localized in cytoplasmic vesicles and as previously described [27], a signal was also detected at the Golgi/Endosome-Like Compartment (ELC) area (Fig 1B). During cytokinesis, at the onset of daughter cell budding, this Golgi/ELC localization of Rab11A was clearly visualized in emerging daughter cells, together with a strong enrichment of the protein at the apical tip of the growing buds, reflecting a possible Rab11A-dependent transport of newly synthesized material between these two locations. Rab11A also accumulates at the basal pole of the parasite both during the G1 phase and cytokinesis (Fig 1B). In order to get further insights into the dynamic localization of Rab11A, we used the previously established transgenic ddFKBP-myc-mCherryRab11A-RHΔ*Ku80* parasites (from here designated as mcherryRab11A-WT parasites) [27] [30]. In this strain, the expression of Rab11A fused to a mCherry tag is under the control of an N-terminal ddFKBP tag, which allows regulation of recombinant protein levels by the inducer Shield-1. Using super-resolution live imaging of parasites expressing the Inner Membrane Complex protein IMC3-YFP and mCherryRab11A-WT, we clearly observed bi-directional trajectories of Rab11A-positive vesicles between the basal and the apical poles of the parasite both, within the parasite cytosol (Fig 1C and Suppl. Movie SM1) and along the parasite cortex delineated by the IMC3-YFP staining (Fig 1C and Suppl. Movie SM2). We also confirmed by videomicroscopy the enrichment of Rab11A at the Golgi/ELC area of newly formed daughter cells and the transport of Rab11A vesicles along the daughter bud scaffold (Suppl. Movie SM3). Interestingly, we also noticed Rab11A-positive dynamic vesicles and tubular structures in the residual body region (Fig 1C, RB). This region has been recently described to harbor a dense actino-myosin network that interconnects the intracellular dividing tachyzoites [6] [7], suggesting that Rab11A may regulate actin-dependent material exchanges between the parasites or the dynamics of this cell-to-cell connecting network. In line with this observation, after transient expression of actin chromobodies coupled to Emerald GFP (Cb-E) that specifically label filamentous actin [7], we visualized Rab11A-positive vesicles moving along actin-positive structures at the parasite cortex (Fig. 1D, upper panel and Suppl. Movie SM4) or anchored to dynamic F-actin structures within the parasite cytosol (Fig. 1D, lower panel and Suppl. Movie SM5). As previously observed [7], we also detected very dynamic F-actin structures at the Golgi/ELC area that co-distribute with the Rab11A signal (Suppl. Movie SM4), suggesting that vesicle budding and/or transport from these compartments may depend on the actin cytoskeleton. To investigate whether Rab11A-positive vesicle movements depend on the actin cytoskeleton, we treated IMC3-YFP / mcherryRab11A-WT tachyzoites with cytochalasin D (CD) for 30 minutes before recording parasites by live imaging. Depolymerizing actin filaments by CD prevented vesicle trafficking and led to the formation of *quasi* static cytosolic and cortical Rab11A-positive clusters (Fig 1E, Suppl. Movie SM6).

**Figure 1.**
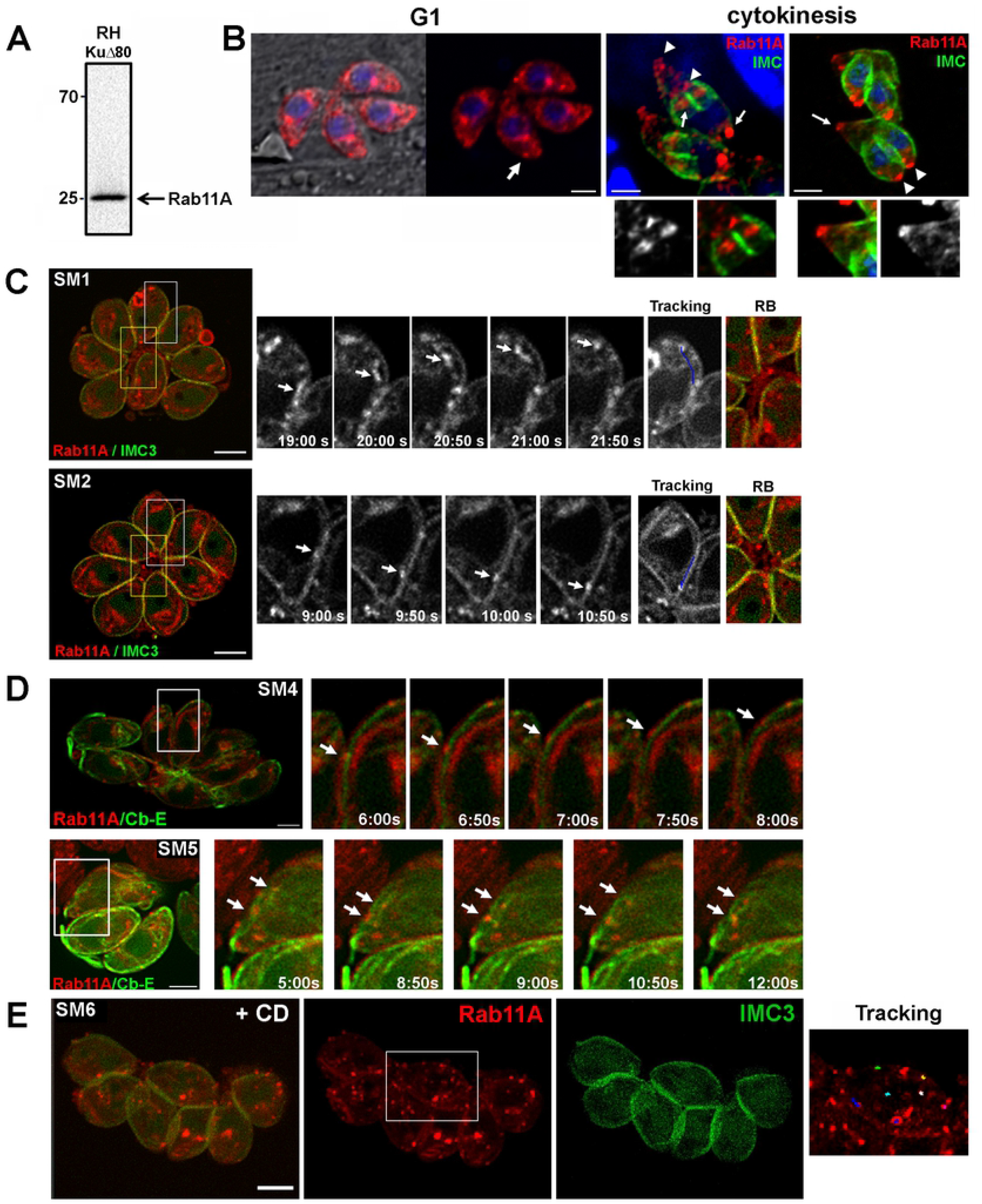
**A-** Western blot analysis with specific anti-Rab11A antibodies detects a unique band at 25kDa in RHΔ*KU80* parasite lysate. **B**-Analysis of Rab11A localization in fixed RHΔ*KU80* parasites using antibodies recognizing Rab11A and IMC3 as indicated. Bars: 1μm. **C**-Sequences of images extracted from movies SM1 and SM2 (left images, white frames) showing the dynamic bi-directional movement of Rab11A-positive vesicles in the cytosol (upper sequence) and along the parasite cortex (lower sequence) of mcherryRab11A-WT and IMC3-YFP expressing parasites. Tracking of vesicle trajectory is also shown. Images on the right show a zoom of the residual body (RB) region indicated by a yellow frame in the corresponding vacuole. Bars: 2 μm. **D-** Sequences of images extracted from movies SM4 and SM5 (left images, white frames) showing the dynamic movement of Rab11A-positive vesicles along the actin-positive parasite cortex (upper sequence) and their interaction with dynamic F-actin structures within the parasite cytosol (lower sequence) of mcherryRab11A-WT and Cb-Emerald expressing parasites. Bars: 2 μm**. E**-Images extracted from movie SM6, where mcherryRab11A-WT and IMC3-YFP expressing parasites were treated with cytochalasin D (CD) for 30 min before being recorded. Rab11A-positive vesicles localized in *quasi* static clusters, as shown after tracking the trajectories. Bar: 2μm.

Collectively, these data demonstrated that Rab11A-positive vesicle movement is dependent on the actin cytoskeleton activity and that Rab11A might participate in (i) vesicle budding from the TGN/ELC, (ii) cargo transport between the apical and basal poles of the parasite and (iii) material exchange between the replicating parasites via release of vesicles at the basal pole.

### Rab11A-positive vesicles dynamically co-distribute with DG

The DG-mediated secretory pathway is considered in *T. gondii* to be the default constitutive secretory pathway based on the observation that the soluble SAG1 protein truncated of its GPI anchor (SAG1ΔGPI) is transported within DGs before being released into the vacuolar space [10] [31]. Interestingly, the dynamic motion of Rab11A-positive vesicles was very similar to the recently described actin and myosin F-dependent movements of DGs [31] and Rab11A is a known regulator of exocytosis in other eukaryotic systems [14].

In order to explore dense granule dynamics in relation to Rab11A, we expressed SAG1ΔGPI-GFP in mcherryRab11A-WT parasites. Using live imaging, we confirmed that the DG content was efficiently released as illustrated by the localization of the GFP signal into the vacuolar space (Fig 2A). The GFP-positive DGs detected in the parasite cytosol displayed a significant and dynamic co-distribution with mcherryRab11A-WT positive vesicles (Fig 2B, 2C). 33,7% of the DG population co-distributed over time with Rab11A-positive vesicles in replicating tachyzoites, while 26,1% of Rab11A-positive vesicles co-distribute with DGs. In agreement, the fluorescent signal intensity profile indicates that GFP-positive DG and mcherryRab11A-positive vesicles are closely apposed (Fig 2A and 2B). This is also clearly visualized in the movie SM7 (Fig 2D), in which a DG appeared docked onto a Rab11A-positive vesicle, the latter being anchored at the periphery of the parasite, and both compartments are simultaneously transported along the parasite cortex (Fig 2D). We tracked this GFP-positive DG motion (Fig 2E and 2F; Suppl. Movies SM8 and SM9) and fitted the recorded xy positions over time using mathematical models of “directed” or “diffusive” motion (see M&M) [32]. We confirmed that the DG trajectory 2 is consistent with a “directed” motion (fitted curve, Fig 2F), characteristic of a vesicle moving along cytoskeleton tracks, in contrast to the trajectories 1 and 3, consistent with “confined diffusive” motions [32]. This result, together with the inhibition of Rab11A-positive vesicle (Fig 1D) and DG [31] movements upon CD treatment, strongly suggests that Rab11A promotes DG transport by mediating DG tethering along actin filaments, at least at the parasite cortex.

**Figure 2.**
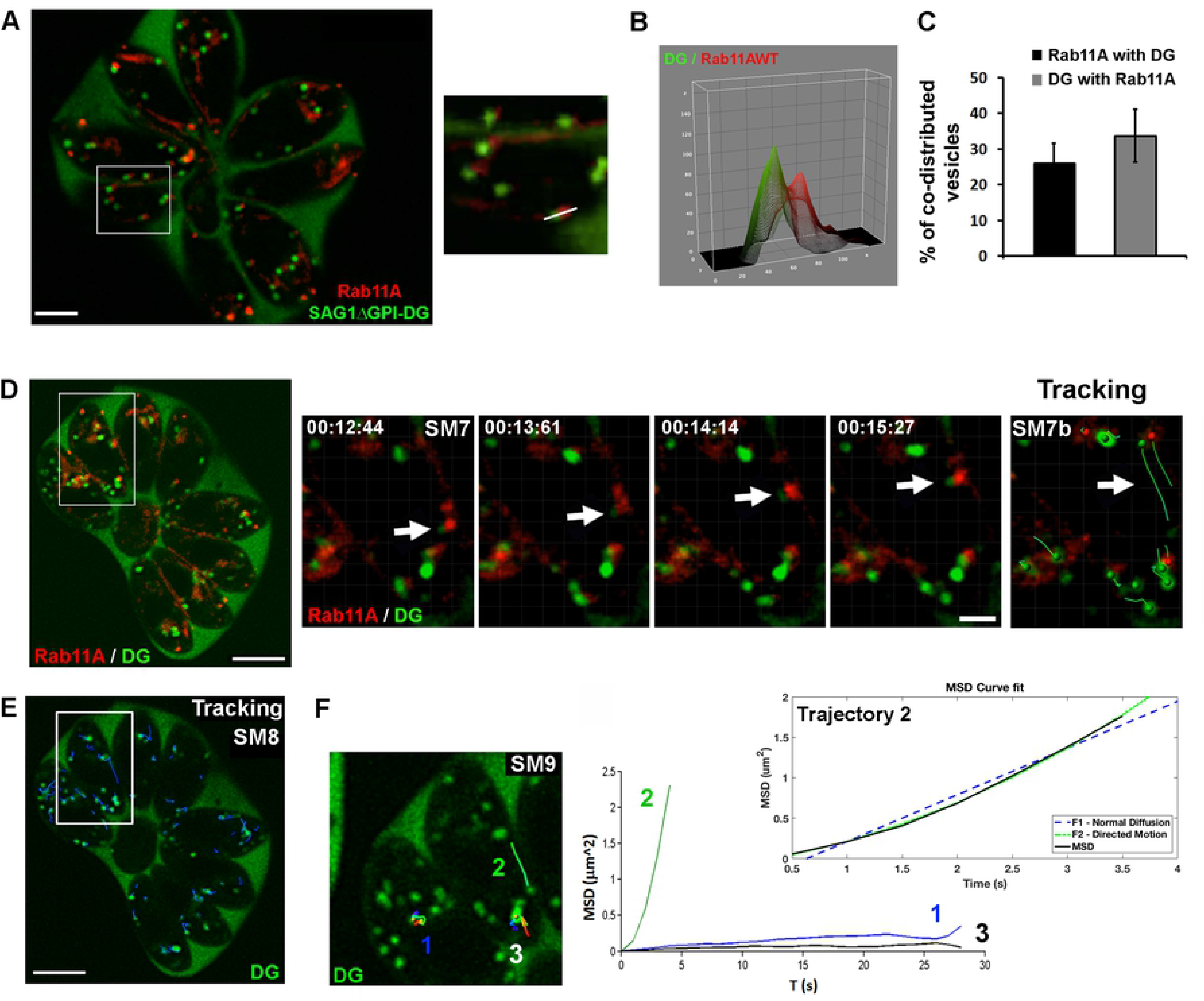
**A-** Image extracted from a time-lapse acquisition illustrating the release of SAGΔGPI protein (green) into the vacuolar space of mcherryRab11A-WT and SAGΔGPI-GFP expressing parasites, as well as the co-distribution in the parasite cytosol of SAGΔGPI-GFP positive DG and mcherryRab11A-WT positive vesicles (red). The right insert shows a zoom of the region indicated by a white frame in the full vacuole. Bar: 2μm. **B**-Fluorescence intensity profiles plotted over the distance of the GFP and mcherry signals along the line indicated in A (insert). **C-** Percentage of co-distribution between the total population of SAGΔGPI-GFP-positive DGs and mcherryRab11A-WT-positive vesicles of a given vacuole averaged over 5 consecutive time points (n=10 vacuoles). Data show mean ± sd. (**p<0,001). **D-** Sequences of images extracted from movie SM7 (region indicated by a white fame in the full vacuole) illustrating the joint motion of a Rab11A-positive vesicle (red) and a SAGΔGPI-positive DG (green) along the parasite cortex, as illustrated by their tracking. Time is indicated in seconds. **E-** Automated tracking of all DG trajectories within the vacuole (SM8). **F-** Three trajectories (1, 2, 3) (Movie SM9) in the region indicated by a white frame in **E**-were analyzed by plotting the Mean Square Displacement (MSD) over ΔT (s) using the Imaris software. Trajectory N°2 (black line) corresponding to the track shown in **-D** (SM7) fitted a mathematical model of “directed” motion (green line) defined by the equation MSD=4Dt+v^2^t^2^ while trajectories 1 and 3 displays a “confined” motions.

### Rab11A promotes DG exocytosis

To assess whether Rab11A regulates DG transport, docking or the later step of fusion at the PM, we used a previously established parasite strain that over-expresses in a rapidly inducible manner an inactive GDP locked version of Rab11A fused to the mCherry fluorescent reporter (DDmCherrycmycRab11A-DN-RHΔ*Ku80*; from here called mCherryRab11A-DN and distinguished from mCherryRab11A-WT) [27] [30]. By WB, we confirmed that both Rab11A-WT and Rab11A-DN proteins were expressed in similar amounts after 4 h induction with Shield-1 (Fig 3A). First, we monitored DG release in fixed Rab11A-WT and Rab11A-DN intracellular tachyzoites following gentle saponin permeabilization, which improved detection of secreted GRA proteins localized in the vacuolar space and at the PVM. To rule out any indirect effect of the previously described cytokinesis defect on DG secretion in Rab11A-DN parasites [27], we pre-treated freshly egressed extracellular tachyzoites for 1 h with Shield-1 before seeding them on a fibroblast monolayer and analyzed DG secretion 2h and 4h after parasite invasion (Fig 3B). Our data revealed a highly significant block of GRA1 and GRA3 secretion in Rab11A-DN parasites in contrast to Rab11A-WT in which both proteins were typically released in the vacuolar space or decorated the PVM (Fig 3B and 3C). A similar observation holds for additional GRA proteins (GRA2, GRA5, GRA6 and GRA16) as shown in Suppl. Fig 1. Notably, in contrast to Rab11A-WT parasites, GRA16-positive DGs were also retained within Rab11A-DN parasite cytosol and accordingly GRA16 no longer reached the host cell nuclei 16h post-infection [33] (Suppl. Fig 1B).

**Figure 3.**
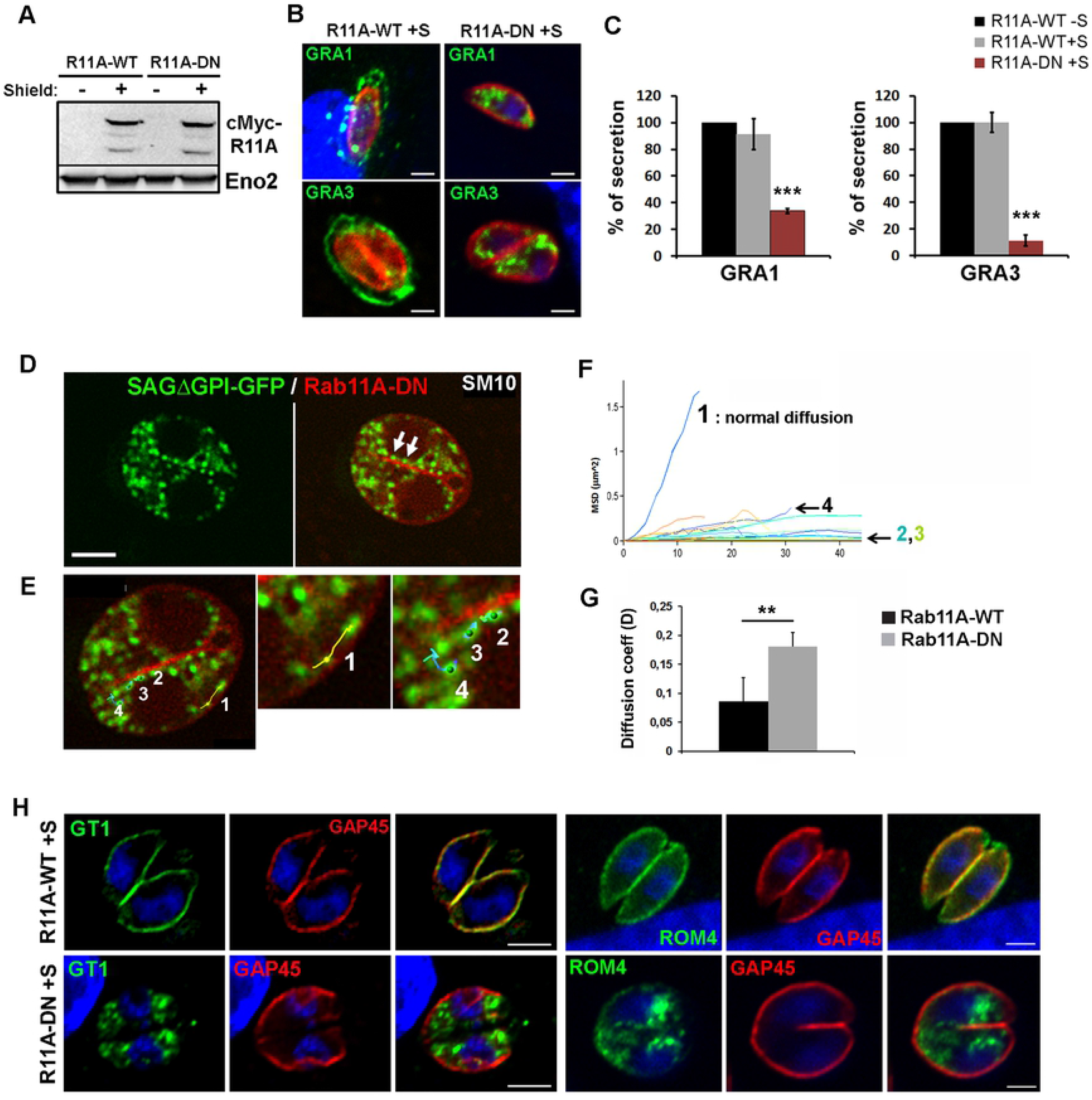
**A-** Western blot analysis with anti-Rab11A antibodies detects Rab11A-WT and Rab11A-DN proteins in similar amounts after 4 h of Shield-1 induction (+S) of intracellular tachyzoites. Eno2 is used as a loading control. **B-** Immunofluorescence assay (IFA) showing the dense granule proteins GRA1 and GRA3 (green) retained in intra-cytosolic vesicles following 2 h (upper panel) and 4 h (lower panel) of Shield-1 induction of Rab11A-DN parasites, while being efficiently released into the vacuolar space and at the vacuole membrane in similarly induced Rab11A-WT parasites. The parasite cortex is delineated by the glideosome protein GAP45 (red). Bars: 1μm. **C-** Percentage of vacuoles positive for GRA1 and GRA3 secretion in Rab11A-WT and Rab11A-DN parasites induced (+S) or not (-S) with Shield-1. Data show mean ± SEM of three independent experiments. **D**-Image extracted from movie SM10 illustrating DG movements in mcherryRab11A-DN / SAGΔGPI-GFP expressing parasites. DGs accumulate in the parasite cytosol and remain stationary along the segregating membrane of daughter cells (arrows). Bar: 2μm. **E-** Images extracted from movie SM10 illustrating the trajectories of 4 DGs analyzed in F-. **F-** Tracking of DGs in Rab11A-DN expressing parasites indicates mostly confined (as exemplified for DG trajectories 2, 3) and diffusive (trajectories 1, 4) motions. **G**-Mean diffusion coefficient (D) calculated from 10 cortical trajectories manually tracked in Shield-1 induced Rab11A-WT and Rab11A-DN parasites. Data show mean ± sd. (**p<0,01). **H-** IFA showing the glucose transporter GT1 and Romboïd protein ROM4 (green) retained in intra-cytosolic vesicles in Shield-1 induced Rab11A-DN parasites, while being efficiently delivered at the plasma membrane in induced Rab11A-WT parasites. The parasite cortex is delineated by GAP45 (red). Bars: 2μm.

To further analyze the role of Rab11A in DG secretion, we also expressed SAG1ΔGPI-GFP in mcherryRab11A-DN parasites. In contrast to Rab11A-WT parasites, Rab11A-DN parasites were drastically impaired in their ability to release SAGΔGPI-GFP into the PV space (Fig 3D). Consequently, DGs were densely packed in the cytosol, which impaired reliable automatic tracking of all vesicles and therefore the quantification of the percentage of directed *versus* diffusive trajectories in the total DG population. Nonetheless, DGs appeared to mostly display normal diffusive and confined motions (Fig 3F and Suppl. Movie SM10). In particular, the accumulation of DGs observed at the altered interface between the two segregating daughter cells accounted for a local *quasi* static behavior as illustrated by their confined trajectories (Fig 3F: trajectories 2, 3 and Suppl. Movie SM10). Few longer-range trajectories could be detected along the cortex of the parasites (such as illustrated for trajectory 1), however they never fitted a model of directed motion with good probability. In support of this result, analysis of cortical DG trajectories in Rab11A-WT and Rab11A-DN parasites revealed a significant increase in the coefficient of diffusion of Rab11A-DN trajectories, suggesting a role for Rab11A in regulating DG directed transport along the parasite cortical cytoskeleton (Fig 3G). Finally, we performed an experiment in which we washed out 0.5μM- (SM11) or 1μM- (SM12) Shield-1 pre-induced Rab11A-DN parasites in order to stop the expression of the Rab11A-DN protein. 4h after Shield-1 removal, we clearly observed a strong accumulation of GFP-positive DGs at the PM separating dividing parasites together with the re-initialization of their content release (Suppl. Movies SM11 and SM12). Since this PM accumulation was not detected in Rab11A-DN parasites in presence of Shield-1, this suggests that Rab11A is required for the early step of DG docking/tethering at the PM. Of note, as vesicle docking/tethering precedes the final fusion step of the exocytic process, we could not decipher whether Rab11A is also involved in the fusogenic process itself.

Collectively, our data indicate that Rab11A regulates both the directed transport of DG along cytoskeleton tracks (Fig 1D and Fig 2D, E, F) and their exocytosis in the PV space likely by promoting DG docking/tethering at the parasite PM.

### Rab11A regulates transmembrane protein localization at the PM

Based on our previous study [27], we proposed that Rab11A is required for the delivery of vesicles containing SAG1 and probably other surface proteins, from the endosomal network to the plasmalemma of daughter cells, where new PM is synthesized, similar to the function described in other eukaryotes. This prompted us to investigate whether Rab11A might regulate the localization of other surface proteins in *T. gondii* during replication. We transiently transfected Rab11A-WT and Rab11A-DN parasites with plasmids encoding the transmembrane HA-tagged Glucose transporter 1 (GT1) [34] or the Ty-tagged rhomboïd protease 4 (ROM4) [35]. In contrast to the rhomboid protease ROM1 that localizes to micronemes, ROM4 was found to be targeted to the tachyzoite PM, suggesting that it may be transported through the constitutive pathway [35] [36]. Similar to DGs, GT1 and ROM4 proteins were retained in intracellular vesicles and were no longer delivered to the parasite PM (Fig 3H). In addition, we took advantage of the impaired exocytosis activity in Rab11A-DN parasites to study whether different populations of secretory vesicles may co-exist during parasite replication. Co-localization studies in fixed parasites showed that ROM4 and GRA3 partially co-localize but were also detected in distinct vesicular compartments both immediately after parasite invasion when de novo synthesis of DG proteins occurs and also after the first division cycle (Suppl. Fig 1C). This may reflect a distinct timing of protein synthesis and vesicle release from the Golgi to the PM but it also suggests the existence of different regulatory pathways for the trafficking of protein localized at the PM *versus* proteins secreted into the vacuolar space. In particular, transmembrane proteins may be actively recycled during parasite division as suggested in a previous study on the retromer subunit TgVPS35 [37] and more recently during extracellular parasite motility [38]. Thus, our data indicate a broader role of Rab11A-mediated exocytosis for the delivery of proteins at the PM and for the release of DG proteins into the vacuolar space during parasite replication.

Importantly, unlike GRA protein secretion, DG biogenesis was not impaired in Rab11A-DN parasites as assessed by transmission electron microscopy (Fig 4). In addition, supporting a major disturbance in DG exocytosis, the IVN could not be detected in the drastically reduced vacuolar space characterized by the PVM being closely apposed to the parasite PM (Fig 4B and 4C). We also detected the previously described defect in daughter cell membrane segregation [27] (Fig 4C, arrows).

**Figure 4.**
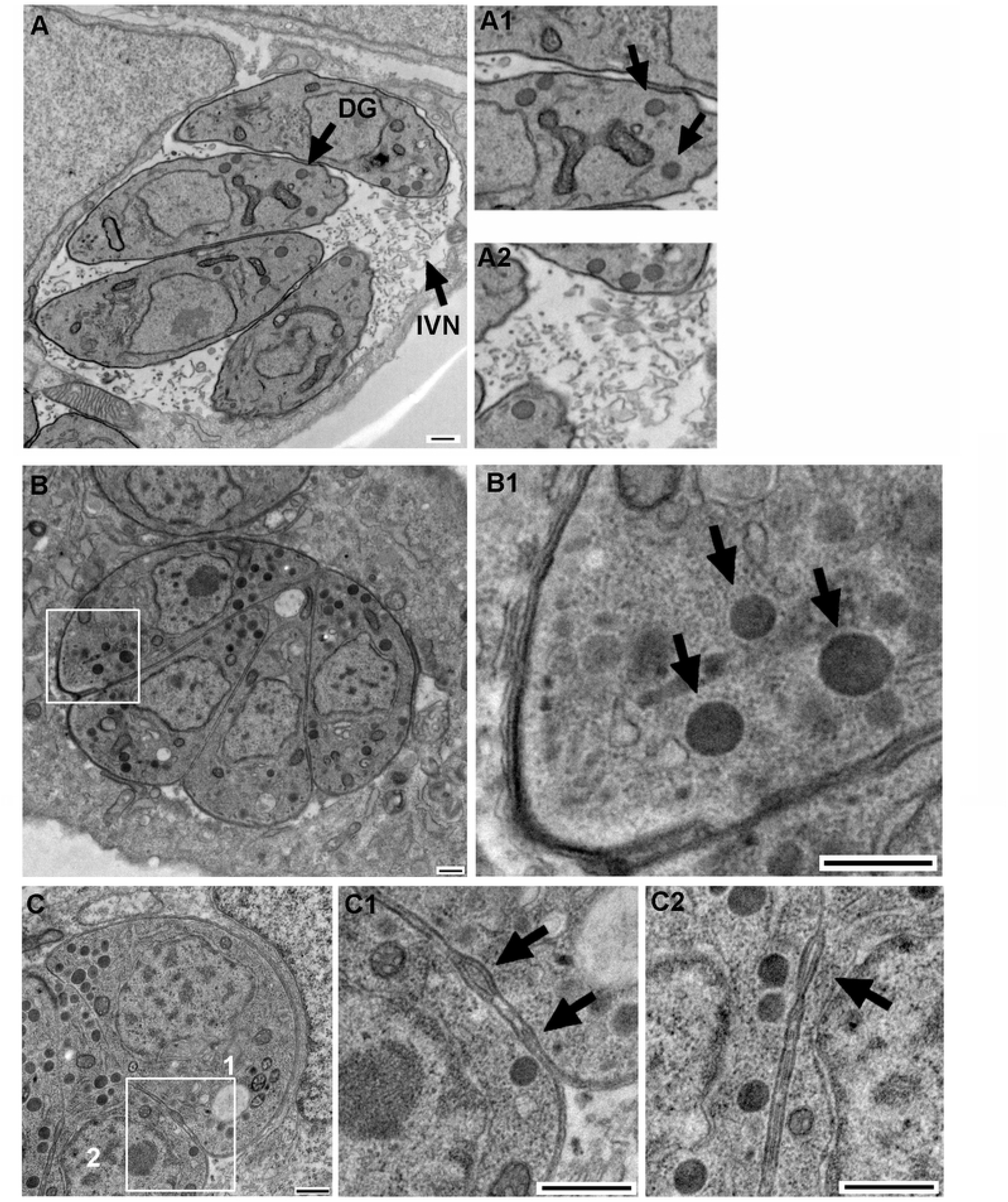
Electron micrographs of infected host cells harboring Shield-1 induced Rab11A-WT replicating parasites (**A**), in which dense granules (A1) and the IVN (A2) are visualized. Shield-1 induced Rab11A-DN parasites (**B**) accumulate dense granules (B1), notably at their basal pole and the IVN is not detected in the drastically reduced vacuolar space. Rab11A-DN expressing parasites also display a previously described defect in membrane segregation between daughter cells (**C**). A zoom of the regions 1 and 2 is shown in C1 and C2. Bars: 500nm.

### Rab11A regulates adhesion and motility of extracellular parasites

A role for Rab11A in parasite invasion has been previously demonstrated [30]. To explore which steps of parasite entry (e.g. adhesion, motility, invasion) were altered, we treated extracellular Rab11A-WT and -DN parasites with Shield-1 for 2 h before monitoring their ability to adhere to host cells. We found that Rab11A-DN tachyzoites were severely impaired in their surface attachment to human fibroblast (HFF) monolayers compared to Rab11A-WT parasites (Fig 5A). Furthermore, parasites that successfully adhered exhibited a strong defect in motility as quantified by the percentage of parasites displaying a SAG1-positive trail deposit (Fig 5B). Importantly, compared to Rab11A-WT parasites, the morphology of adherent motile Rab11A-DN parasites was altered, the latter being wider and shorter, losing their typical arc shape (Fig 5C). Analysis of individual parasites imaged by Scanning EM (n=70 for WT and DN) confirmed that Rab11A-DN parasites display a significant increase in their circularity and accordingly, a decrease in their aspect ratio (AR: major axis/minor axis) (Fig 5C). However, the apical conoid with the emerging microtubule array could be visualized, suggesting no defect in the establishment of parasite polarity (Fig 5C). An impaired recruitment of late glideosome components at the daughter cell buds has been previously reported in dividing Rab11A-DN parasites [27] and could account for the motility defect. However, we induced Rab11A-DN protein expression in non–dividing extracellular parasites and accordingly we did not observe any significant defect in the localization of GAP45 and Myosin Light Chain 1 (MLC1) at the parasite cortex of extracellular parasites (Suppl. Fig 2). This indicates that the morphological defect observed in Rab11A-DN parasites is not correlated with a significant perturbation of glideosome component localization.

**Figure 5.**
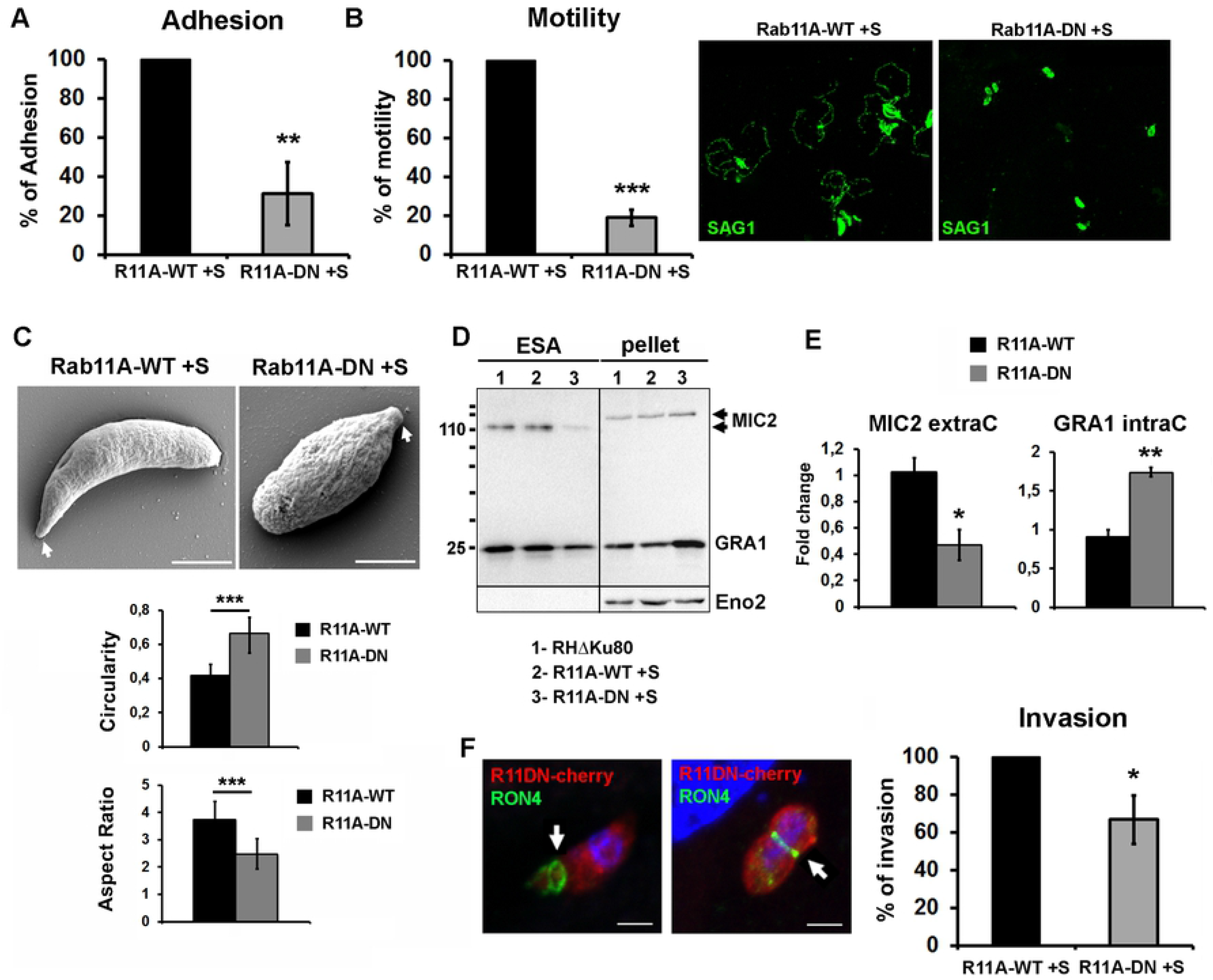
**A-** Quantification of the percentage of Shield-1 induced extracellular Rab11A-DN parasites adhering to host cells normalized to control Rab11A-WT parasites. Data show mean ± SEM of three independent experiments. (**p<0,01). **B-** Quantification of the percentage of Shield-1 induced extracellular Rab11A-DN parasites, normalized to control Rab11A-WT parasites, displaying a SAG1-positive trail deposit (green) as illustrated in the images. Data show mean ± SEM of three independent experiments. (***p<0,001). **C-** Scanning Electron Micrographs (SEM) of Shield-1 induced extracellular Rab11A-WT and Rab11A-DN parasites, which were allowed to move for 15 min on BSA-coated coverslips before fixation. Arrows indicate the apical pole of the parasite. Bars: 2μm. The histograms indicated the mean Circularity (upper graph) and Aspect Ratio (major axis / minor axis) (lower graph) of Shield-1 induced extracellular Rab11A-WT and Rab11A-DN parasites imaged by SEM (n=70 parasites for each condition; ***p<0,001). **D-** Western blot analysis of excreted-secreted antigen assays (ESA) performed with Shield-1 induced (+S) or not (-S) extracellular Rab11A-WT and Rab11A-DN expressing parasites revealed a defect in MIC2 and GRA1 protein secretion. Eno2 was used as a loading control. **E-** Quantification of secreted MIC2 proteins (ESA fraction) and intracellular GRA1 proteins (pellet fraction) from 3 independent ESA (as shown in **D**-) expressed in fold-change compared to non-induced Rab11A-WT parasites (lanes 1 in D-) (*p<0,05, **p<0,01). **F-** Quantification of the percentage of Shield-1 induced extracellular Rab11A-DN expressing parasites, which have invaded host cells normalized to control Rab11A-WT expressing parasites. Data show mean ± SEM of three independent experiments. (*p<0,05). Fluorescence images show Shield-1 induced mcherryRab11A-DN (red) invading host cells, as illustrated by the presence of a circular junction positive for RON4 (green). Bars: 1μm.

The microneme protein MIC2, a transmembrane protein released at the PM of the parasite, promotes parasite adhesion and motility [39] [40]. First, we confirmed by IFA that MIC2-positive micronemes were detected at the apical pole of extracellular induced Rab11A-DN parasites, indicating no major defect in their localization (Suppl. Fig 3A). Secretion of microneme proteins by extracellular parasites can be triggered by ethanol, a step followed by their release from the parasite PM after cleavage by proteases. Notably, ROM4 has been shown to promote MIC2 trimming at the parasite PM. Since ROM4 was no longer present at the PM of replicative Rab11A-DN parasites, we investigated whether a similar defect could be observed in 2h Shield-1 induced extracellular Rab11A-DN. As previously observed for glideosome components, ROM4 localization at the PM was not perturbed in extracellular induced Rab11A-DN parasites (Suppl. Fig 3B). Next, we performed excretion/secretion assays to assess the transport of MIC2 protein to the parasite PM and its subsequent release in the culture medium. Western blot quantification of the Excreted-Secreted Antigen (ESA) fractions demonstrated a significant reduction in MIC2 release upon induction of microneme exocytosis. Accordingly, a slight increase in MIC2 protein level was observed in the pellet fraction, also indicating that the observed decrease in MIC2 secretion is not due to a defect in protein synthesis. As observed by IFA, a reduced level of constitutive GRA1 secretion was also detected by WB, which correlates with GRA1 accumulation in the parasite pellet fraction (Fig 5D and 5E). Thus, our data suggest that the defect of extracellular Rab11A-DN parasites in host cell adhesion and motility MIC2 are at least partially due to an impairment of efficient MIC2 delivery to the PM.

Lastly, Rab11A-DN parasites that successfully adhered to the surface of host cells, displayed only a mild defect in host cell invasion (Fig 5F). This was supported by the observation of a correctly formed RON4-positive junction by invading Rab11A-DN parasites (Fig 5F).

Collectively, our results demonstrate that Rab11A promotes parasite invasion by regulating parasite adhesion and motility, but not the formation of the circular junction. This defect correlates with severe morphological alterations of extracellular parasites.

### Rab11A-positive vesicles accumulate at the apical pole during parasite motility and host cell invasion

The active role of Rab11A in parasite adhesion and motility led us to explore the localization of Rab11A in motile extracellular and invading parasites. Live imaging of mcherryRab11A-WT revealed an unexpected polarized motion of Rab11A-positive vesicles towards two main foci localized at the apical tip of extracellular adhering and motile parasites (Fig 6A, Suppl. Movie SM13). This process appeared to be prolonged during host cell invasion (Fig 6B, SM14). This Rab11A apically polarized localization in invading parasites was further confirmed in fixed parasites after labeling of the circular junction using the RON4 marker (Fig 6C).

**Figure 6.**
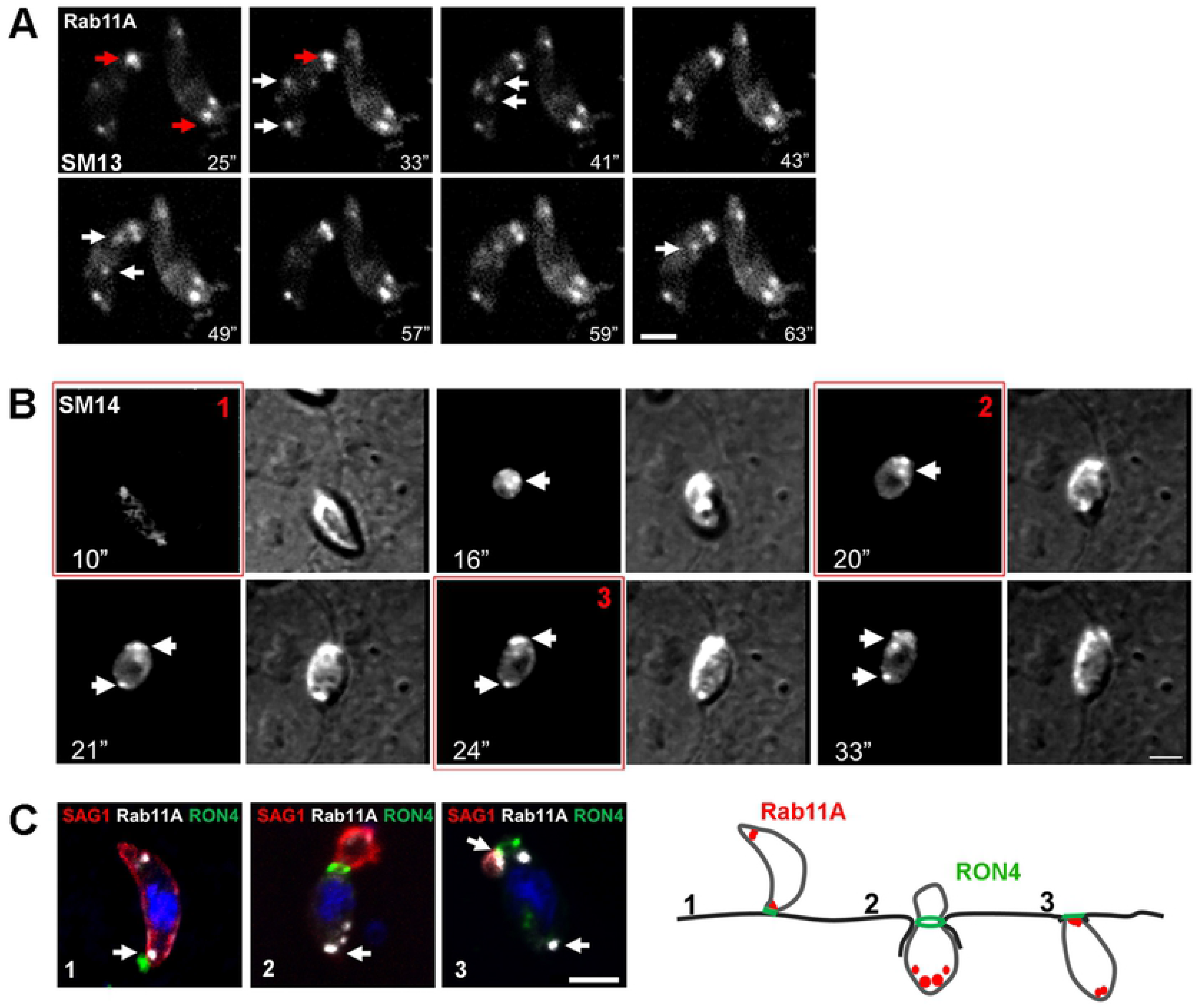
**A-** Sequences of images extracted from movie SM13 showing the polarized recruitment of mcherryRab11A-positive vesicles (white arrows) towards two main foci localized at the tip of adhering parasites (red arrows). Time is indicated in seconds. Bar: 2μm. **B-** Sequences of images extracted from movie SM14 showing a similar polarized localization of mcherryRab11A-positive vesicles (white arrows) during host cell invasion. Time is indicated in seconds. Bar: 2μm. **C-**Fluorescence images of RHΔ*KU80* parasites fixed at three different steps of the host cell invasion process, as illustrated in the right scheme. The circular junction is labeled with RON4 (green) and the membrane protein SAG1 was used to label the extracellular portion of the invading parasite (red). Bar: 2μm.

### Rab11A regulates polarized secretion of DG content during parasite motility and host cell invasion

Next, we assessed whether Rab11A regulates the secretion of DGs, not only during parasite replication (Fig 3) but also during parasite motility and invasion. Similarly to our live imaging data (Fig 6), we found a clear localization of Rab11A at two foci localized at the apex of extracellular parasites that have been allowed to move on coverslips prior fixation (Fig 7A). These Rab11A foci strongly co-localized with the DG protein GRA1, suggesting that the apically polarized secretion of DGs may play a role in the regulation of parasite adhesion and motility. A similar co-recruitment of both, Rab11A and DG at two apical foci was observed during host cell invasion (Fig 7B). Most importantly, we observed a complete inhibition of this polarized secretion of DG in extracellular motile induced Rab11A-DN parasites (Fig 7A) and during host cell invasion (Fig 7B). This result demonstrates that Rab11A regulates the apically polarized secretion of DGs during the early steps of parasite adhesion and entry into host cells. This apical DG secretion may reflect the delivery of a new membrane pool or regulatory factors contributing to parasite motility.

**Figure 7.**
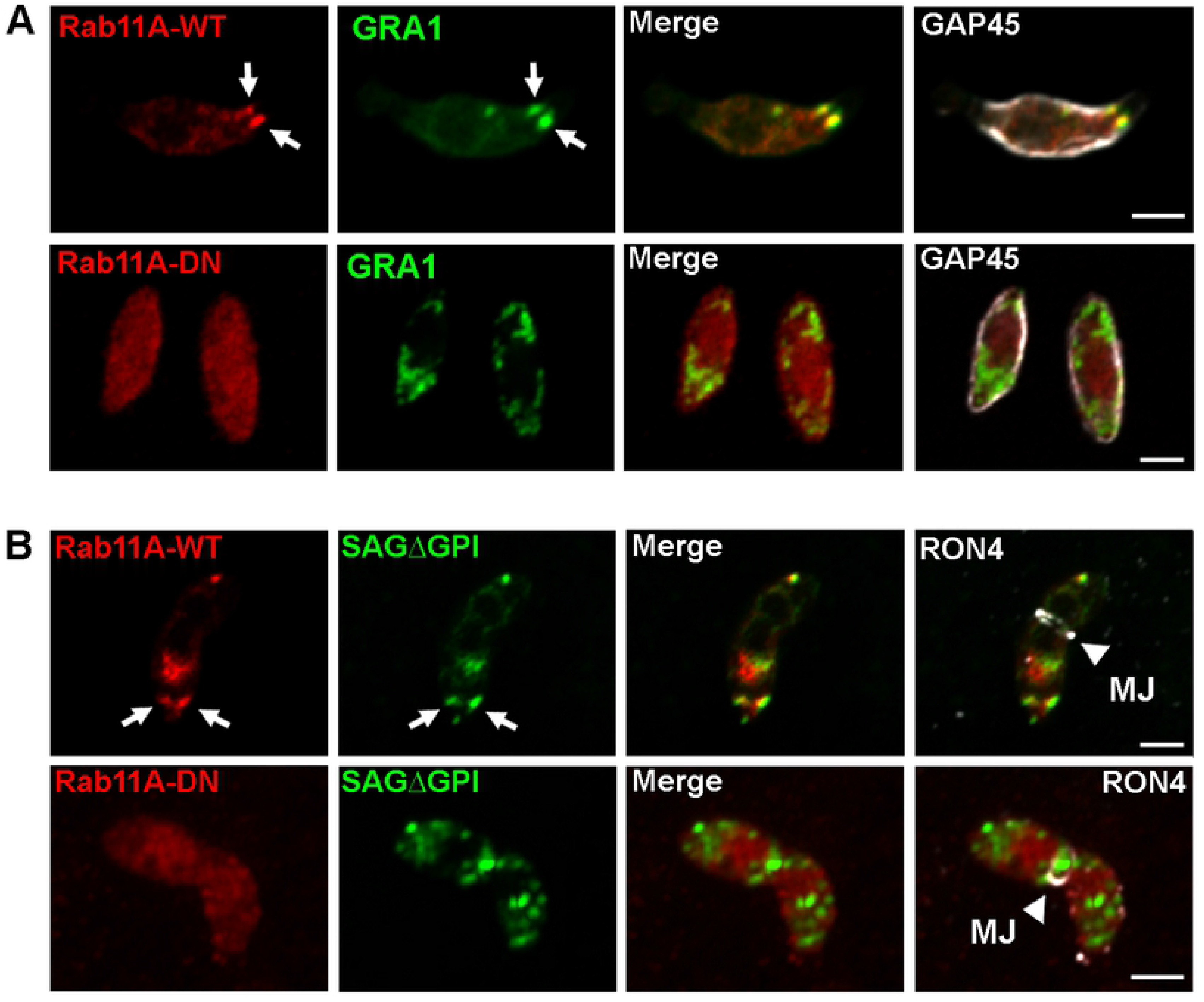
**A-** Immunofluorescence images showing the co-localization of the mcherryRab11A-positive signal (red) and GRA1-positive DG at two apical foci localized, where the Inner Membrane Complex (labeled with anti-GAP45 antibodies, white) interrupts, in motile extracellular induced Rab11A-WT (upper raw). This apically polarized secretion is no longer detected in induced Rab11A-DN expressing parasites (lower panel). Bars: 2μm. **B-** A similar apical and focalized co-localization between Rab11A and SAGΔGPI-GFP-positive DGs is observed during host cell invasion (illustrated by the detection of the RON4-positive circular junction). DG apical secretion is no longer observed in invading Rab11A-DN. Bars: 2μm.

## Discussion

In this study, we unraveled an essential role of Rab11A in the delivery of transmembrane proteins at the parasite PM and the release of DG proteins into the vacuolar space.

In other eukaryotic systems, Rab11A localizes to the endocytic recycling compartment (ERC) and has been implicated in the trafficking of internalized receptors from the ERC to the PM [8]. Rab11A also localizes to the TGN compartment, where it regulates transport of material from this compartment to the ERC or the PM [16]. Similarly, during *T. gondii* cytokinesis, Rab11A localizes to the Golgi/ELC region of daughter cells, and at the tip of growing buds, suggesting a polarized transport of *de novo* synthetized material between these two locations during daughter cell emergence. Interestingly, such apically polarized localization of Rab11A was also evident during extracellular parasite adhesion and motility. Thus, one may envision that components of the apical complex, a microtubule-rich structure from which emanates the subpellicular microtubules [41], may control Rab11A-dependent recruitment and exocytosis of specific cargos at the apical pole of the parasite. In particular, RING2, a component of the apical polar ring, was shown to function in constitutive and cGMP-stimulated secretion of microneme proteins [42]. More recently, two other components of the apical polar ring, APR1 and the Kinesin A, were also reported to regulate MIC2 secretion [42]. Hence, it will be of interest to investigate whether Rab11A could also interact with components of the apical polar ring to promote exocytosis of micronemes and DG content during extracellular motility.

Moreover, during the G1 phase of the cell cycle, videomicroscopy recordings of mCherryRab11A-WT expressing parasites revealed highly dynamic Rab11A-positive vesicles displaying bidirectional trajectories between the apical and the basal poles, with a pronounced accumulation at the basal pole of the parasite, consistent with this location being a preferential site for exocytic events. In line with this, the double membrane of the IMC may be considered a major physical barrier for internalization and secretion of material at the parasite plasma membrane. Therefore, it is possible that exocytic events may be enhanced at sites where the IMC interrupts, e.g. at the apical and basal ends of the parasite. In agreement, previous studies showed massive exocytosis of the DG protein GRA2 in multi-lamellar vesicles at the basal pole of the parasite shortly after entry [43]. We also observed Rab11A-positive vesicles and tubular structures in the region of the residual body, which interconnects parasites during intracellular replication. This region was recently reported to harbor a dense actin-myosin network that connects the parasites within the PV ensuring synchronous divisions [6] [7]. Thus, Rab11A- and actin-dependent vesicular transport may regulate exchanges between parasites within the vacuole. Alternatively, Rab11A may also contribute to the regulation of the actin network function and dynamics. Indeed, in plants, it has been shown that dysregulated Rab11A activity affects actin organization in the apical region of growing pollen tubes [44]. Supporting the hypothesis of a specific interaction between Rab11A and the actino-myosin cytoskeleton in vesicle transport, depolymezing actin filaments results in the formation of *quasi* static cytoplasmic and cortical Rab11A-positive vesicle clusters. A role for the complex Myosin Vb-FIP2-Rab11A in promoting actin-mediated transport of vesicles has been previously observed in mammalian cells [45] [46] [47]. So far, no homologues of Rab11-family interacting proteins (FIPs) have been identified in *T. gondii* and *Plasmodium*. Nonetheless, *P. falciparum* Rab11A was found to directly interact with the myosin light chain 1 (MLC1/MTIP), which therefore links Rab11A-mediated vesicular transport to unconventional myosins and the actin cytoskeleton [27]. In line with this, over-expression in *T. gondii* of a dominant negative form of myosin A led to similar defects in the completion of cytokinesis, as found when Rab11A-DN is over-expressed [27]. However, as actin depolymerization resulted in the formation of both cytosolic and peripheral Rab11A-positive static vesicles, it is possible that distinct myosins regulate different steps of Rab11A/DG transport e.g. MyoF in the cytosol and from the TGN [31], MyoA at the parasite cortex where the glideosome is located [27], and MyoJ in the cell-to-cell connecting network [6]. Further studies using parasite strains deleted for these molecular motors will address this question.

Moreover, co-distribution studies indicated that Rab11A-positive vesicles associate with dense granules in a dynamic manner. However, we did not observe Rab11A at the limiting membrane of DG. Rather, these two compartments appear to transiently dock one with each other enabling joint transient motions that were particularly evident at the cortex of the parasite. Indeed, tracking of the trajectories of Rab11A-positive vesicles and DG revealed that Rab11A-positive vesicles promoted DG anchoring at the parasite cortex and their rapid “directed” transport. This mode of transport called “hitchhiking” has been recently described in different cell types and has emerged as a novel mechanism to control organelle movement [48]. During this process, the “hitchhiker” benefits from distinct molecular motors present at the surface of the “vehicle”. This mode of transport may have additional advantages for the hitchhiker. Notably, endosomes represent multifunctional platforms that receive specific signals and could drive the transport of hitchhiker cargo to particular regions of the cell. Also, co-movement of cargo may facilitate interactions at membrane contact sites important for organelle maturation, fusion and/or material exchange. Related to this last aspect, we found that over-expression of Rab11A-DN led to a complete block in DG secretion, which indicates an additional role of Rab11A in vesicle tethering at- and possibly fusion with- the PM. Rab11A is known to promote vesicle tethering and fusion at the PM via its interaction with the exocyst complex in other Eukaryotic systems [14]. However, homologues of the different exocyst complex subunits could not be identified *in T. gondii* [49]. Thus, unexplored mechanisms of Rab11A-mediated vesicle fusion likely exist in *T. gondii*.

Benefiting from the fast and efficient induction of the Rab11A-DN protein expression in extracellular parasites, we confirmed the previously described defect in host cell invasion [29]. Of note, our numerous attempts to generate parasites expressing C-terminal tagged Rab11A failed, and notably, our attempts to apply the fast inducible AID knock-down system also failed [50]. This is likely due to the fact that the C-terminal domain of the Rabs contains one or two cysteines recognized by geranylgeranyl-transferases to induce their isoprenylation, a modification required for their association with vesicle membranes. The impaired cell invasion of Rab11A-DN expressing parasites results from a strong defect in parasite adhesion to host cells. Indeed, parasites that successfully adhered to host cells were only mildly perturbed in host cell entry. In addition, the secretion of MIC2, an adhesin essential for parasite adhesion and motility was reduced upon dysregulation of Rab11A activity. Secretion of the GPI-anchored protein SAG1 is also altered in Rab11A-DN expressing parasites [27]. Thus, it is likely that the altered secretion of these two host cell adhesins contributes to the decrease in adhesion and motility of Rab11A-DN parasites. In line with a role of Rab11A in the regulation of surface protein trafficking, we also found a strong defect in the localization of the romboïd protease ROM4 and the glucose transporter GT1 at the PM, indicating a broader role of Rab11A in exocytosis. It is likely that distinct exocytic pathways exist in *T. gondii*, such as described in other organisms. In particular, whether a distinct endosome recycling compartment is present in *T. gondii* requires further exploration. Previous studies highlighted that *T. gondi* has functionally repurposed its endocytic system to serve the secretory pathway of this fast replicating intracellular parasite [51] [52]. In this context, the TGN appears to be a hybrid compartment to which the endosomal markers (Rab5 and Rab7) are tightly associated [52]. Therefore, one may envision that material internalized from the PM reaches this hydrid TGN/ELC compartment before being re-directed to other target membranes, such as the rhoptries, the PM and the degradative vacuole (VAC). Such a recycling process has been recently observed during extracellular parasite motility [38]. Recycling of mother material during daughter cell emergence may also follow this indirect secretory pathway, while *de novo* synthetized proteins may traffic via the direct TGN to PM route.

Finally, exocytosis of DG proteins in *T. gondii* is commonly named the “constitutive secretory pathway” due to continuous release of cargos into the vacuolar space during intracellular replication. However, during extracellular parasite motility and invasion, imaging of both live and fixed parasites revealed an unexpected polarized transport of Rab11A-positive vesicles towards two main foci located just beneath the conoïd, where the IMC interrupts. This is consistent with a mechanism of “regulated” secretion triggered upon parasite adhesion to host cells. In mammalian cells, Rab11A-dependent polarized secretion towards the leading edge of motile cells is essential to promote persistent migration [53]. This process provides additional membrane ensuring the extension of the leading edge but also contributes to the translocation of regulators of several signaling pathways, including ones involved in actin and microtubule cytoskeleton activity. In *T. gondii*, the apical exocytosis of some effectors may regulate not only actin-based parasite adhesion and motility but also the modulation of the conoid activity involved in microtubule-dependent motility. Such regulation has been demonstrated for the lysine methyltransferase, AKMT (Apical complex lysine (K) methyltransferase) [54] [55]. Thus, future research will aim to identify the cargos that are apically secreted in a Rab11A-dependent manner and their putative role in regulating parasite motility. We found that some of these factors are likely to be contained in DG as the latter co-localized with Rab11A at the two main apical foci that we observed. Interestingly, it has been recently shown that the DG protein GRA8 contributes to the regulation of parasite motility by regulating conoid extrusion and organization of the microtubule network [56].

Therefore, identifying Rab11A interactors will be an important future goal, as it will improve our understanding of the mechanisms regulating the distinct exocytic pathways in *T. gondii*. In particular, it will be important to characterize the molecular mechanisms involved in Rab11A-positive vesicle anchoring to actin or microtubule molecular motors, and of vesicle tethering and fusion with the PM both, during parasite motility and intracellular replication. Finally, exploring a putative functional interaction between Rab11A-dependent secretion and the apical complex may lead to the discovery of novel regulated secretory mechanisms essential to ensure parasite virulence.

## Materials and Methods

### Parasite culture and transfection

*Toxoplasma gondii* Type I RHΔKU80ΔHXGPRT parasites were grown on confluent Human Foreskin Fibroblast (HFF) cells (CCD-1112Sk (ATCC, CRL-2429^TM^)) which were cultured in complete DMEM (gibcoLife Technologies) supplemented with 10% Fetal Bovine Serum (GibcoLife Technologies) and 1% Pen Strep (gibcoLife Technologies). To obtain the transgenic parasites, the RHΔKU80ΔHXGPRT parental strain was transfected by electroporation following standard procedures with 50μg of the following plasmids.

| Plasmid | Selection | Laboratory |
| --- | --- | --- |
| DD-cmycmcherry-Rab11A-WT | HXGPRT | Meissner M [26] |
| DD-cmycmcherry-Rab11A-DN | HXGPRT | Meissner M [26] |
| IMC3-YFP | DHFR | Gubbels MJ |
| SAG1ΔGPI-GFP | CAT | Heaslip A [31] |
| pLic GRA16-HA | DHFR | MA Hakimi [33] |

Following transfection, parasites were subjected to drug selection and verified for the transfection efficiency by immunofluorescence analysis. Subsequently the parasites were subjected to cloning by serial dilution.

### Production of the anti-TgRab11A antibodies

Recombinant purified GST-Rab11A protein was used to raise a *Tg*Rab11A specific mouse polyclonal antibody. The cleavage site present between the GST tag and Rab11A was digested with Precision protease (GE life science). GST-Rab11A bound to agarose beads was washed with 10 bed volumes of Cleavage buffer (50mM Tris HCl, pH7.0, 150mM NaCl, 1mM EDTA, 1mM DTT) at 4°C. Precision protease (40 units) was added to the cleavage buffer and incubated with the beads at 4°C overnight. The purified Rab11A was collected in the supernatant. 50μg of the purified recombinant protein suspended in Freund’s Adjuvant were injected intra-peritoneally into mice over a series of 4 boosts. Following the third boost, a sample of serum was collected and tested by western blot for antibody reactivity using a total protein extract of parasites. Once specific antibody activity was detected mice were sacrificed and serum collected and stored at −20°C.

### Protein sample preparation and Western Blot

Parasites were lysed in lysis buffer (NaCl 150mM, TrisHCl 20mM, EDTA 1mM, 1% TritonX100, protease inhibitors) and total proteins were subjected to electrophoresis in a 10% polyacrylamide gel. The proteins were transferred onto a nitrocellulose membrane (Amersham^TM^Protran^TM^ 0.45μ NC) by a standard western blot procedure. The membrane was blocked with 5% milk (non-fat milk powder dissolved in TNT buffer: 100mM Tris pH8.0, 150mM NaCl and 0.1% Tween20) and probed with the indicated primary antibodies followed by species-specific secondary antibodies conjugated with HRP. The probed nitrocellulose membranes were visualized using the ECL Western blotting substrate (Pierce).

### Immunofluorescence assay

Confluent HFF monolayers were grown on coverslips and infected with parasites prior to fixing with 4 % PFA for 15 min. After quenching with 50mM NH_4_Cl, the coverslips were permeabilized with 0.2% triton dissolved in 5% FBS-PBS for 30 min. Coverslips were incubated with primary antibodies in 0.1% triton dissolved in 2% FBS-PBS and then washed thrice with 1X PBS. Alternatively, the coverslips were incubated with primary antibodies in 0.01% Saponin diluted in 2%FBS-PBS for 1 h. Incubation with secondary antibodies was performed in 0.1% triton or 0.01% Saponin dissolved in 2%FBS-PBS for 30 min. To label invading parasites, freshly egressed extracellular parasites expressing Rab11A-WT and Rab11A-DN were induced with Shield-1 for 2 h and seeded onto HFF monolayers in a 24-well plate at a concentration of 2*10^6^ parasites (Rab11A-WT) and 4*10^6^ (Rab11A-DN) /500µl complete medium containing Shield-1 per coverslip. The plate was centrifuged for 2 min at 1000rpm at room temperature to trigger adhesion and synchronized invasion events. The plate was immediately shifted to a water bath at 37°C and the parasites were fixed with 4% PFA-sucrose at the following time points - 0, 2 and 5 min. Coverslips were washed with PBS and adherent or invading parasites labeled without permeabilization with the anti-SAG1 antibody and a secondary anti-mouse AlexaFluor405 antibody. After washing with PBS, parasites were permeabilized with 0.05% saponin for 10 min, followed by a blocking step with 5% FBS-PBS for 30 min. Next, coverslips were incubated with rabbit anti-RON4 antibodies and secondary anti-rabbit AlexaFluor488 to label the circular junction. Depending on the experiment, additional primary antibodies were added to detect GRA1, GRA3 and TgRab11A during parasite invasion. Images were acquired using a Zeiss LSM880 confocal microscope equipped with an airyscan module. Antibodies used for IFA experiments were: rabbit anti-HA (Cell Signaling Technology), rat anti-cMyc (Abcam), mouse anti-SAG1 (our lab), rabbit anti-GAP45 (D. Soldati-Favre), mouse anti-MIC2 (V. Carruthers), mouse anti-ROP 2–4 (JF. Dubremetz), mouse anti-GRA2 (Biotem), mouse anti-GRA5 (Biotem), mouse anti-GRA1 (Biotem), rabbit anti-GRA3 (JF. Dubremetz), rabbit anti-IMC3 (MJ Gubbels), rabbit anti-RON4 (M. Lebrun), mouse anti-TY (D. Soldati-Favre) and mouse anti-Rab11A (this study).

### Invasion assay

Freshly egressed extracellular parasites expressing Rab11A-WT and Rab11A-DN were harvested and treated for 2 h with 1µM of Shield-1. Induced parasites were counted and seeded onto HFF monolayers in a 24-well plate at a concentration of 2*10^6^ parasites (Rab11A-WT) or 4*10^6^ parasites (Rab11A-DN) / 500µl complete medium containing Shield-1 / coverslip. The plate was centrifuged for 2 min at 1000rpm at RT to trigger immediate adhesion and synchronized invasion events. Parasites were then shifted to 37°C for 45min. The slips were washed with PBS – three times prior to fixation. Cells were fixed in 4% PFA for 10 min and subjected to a red/green invasion assay. Briefly, adherent external parasites were labeled without permeabilization with mouse anti-TgSAG1 antibodies, followed by secondary anti-mouse antibodies coupled to Alexa488. After cell permeabilization with Triton 0.1%, invaded intracellular parasites were detected using rabbit anti-TgGAP45 antibodies followed with a secondary anti-rabbit antibodies coupled to Alexa594. All parasites labeled green-red were considered as extracellular, while parasites exclusively red (positive for GAP45) were considered intracellular. At least 300 parasites were counted for each condition performed in triplicate. Data represent mean values ± SEM from three independent biological experiments.

### Motility (Trail deposition) Assay

Glass slides were coated with 100µg/ml BSA-PBS and incubated at 37°C for 1 h. The slides were washed three times with PBS and allowed to dry. Freshly egressed extracellular Rab11A-WT and Rab11A-DN expressing parasites were harvested and treated for 2 h with 1µM of Shield-1. Induced parasites were counted and suspended in HHE buffer (HBSS, 10mM HEPES, 1mM EGTA) containing 1µM of Shield-1. 1*10^6^ (Rab11A-WT) or 2*10^6^ (Rab11A-DN) parasites were seeded per well and incubated for 15 min at 37°C. Parasites were then fixed with 4% PFA in PBS for 10 min at RT. A standard IFA protocol was followed wherein primary mouse anti-SAG1 antibodies were used followed by goat anti-mouse secondary antibodies conjugated to Alexa Fluor 488. 200 parasites per coverslip were counted for the presence or absence of a SAG1-positive trail. With internal triplicates, the experiment was performed 3-times. Mean values ± SEM was calculated.

### Adhesion assay

Freshly egressed extracellular Rab11A-WT and Rab11A-DN parasites were harvested and treated for 2 h with 1µM of Shield-1. Parasites were then counted and resuspended in Endo buffer (44.7mM K_2_SO_4_, 10mM Mg_2_SO_4_, 100mM sucrose, 5mM glucose, 20mM Tris, 0.35% wt/vol BSA - pH 8.2) containing 1µM cytochalasin D and 1µM of Shield-1. 2*10^6^ parasites were then seeded onto confluent HFF cells grown on glass coverslips, spun down for 2 min at 1000rpm and incubated for 15 min at 37°C in the presence of 1 μM Cytochalasin D and Shield-1. The coverslips were washed with PBS before fixation with PFA 4% for 10 min. The Red/Green assay was performed (see “Invasion assay”). Data were compiled from 3 independent experiments after counting 20 fields /coverslip at 60X magnification (done in triplicate for each condition/ experiment). Data collected are mean values ± SEM.

### Excreted Secreted Antigens assay

50*10^6^ freshly egressed extracellular Rab11A-WT and Rab11A-DN parasites were harvested and treated for 2 h with 1µM of Shield-1. Shield-1 treatment was maintained throughout the experiment in all media. Parasites were mixed with an equal volume of pre-warmed intracellular (IC) buffer (5 mM NaCl, 142 mM KCl, 1 mM MgCl2, 2mM EGTA, 5.6 mM glucose and 25 mM HEPES, pH 7.2) and spun down at 1500rpm, RT for 10 min. The pellet was washed once in the IC buffer under similar conditions and then resuspended in Egress buffer (142 mM NaCl, 5mM KCl, 1 mM MgCl2, 1mM CaCl2, 5.6 mM glucose and 25 mM HEPES, pH 7.2) containing -/+ 2% ethanol and incubated for 30 min at 37°C. The samples were spun down at 14000 rpm for 15 min at 4°C and the supernatant containing ESA saved. Pellets were washed once in 1x PBS and saved. The ESA and pellet fractions were suspended in 4x Laemelli blue buffer and subjected to Western blot as described above. The blots were probed with mouse anti-MIC2 (V. Carruthers), mouse anti-GRA1 (Biotem) and rabbit anti-eno2 (our lab) antibodies.

### Transmission electron microscopy (TEM)

After infection of a confluent HFF monolayer, cells containing replicating shield-1 induced Rab11A-WT and Rab11A-DN expressing parasites were detached with a scraper, spun down and fixed with 1% glutaraldehyde in 0.1 M sodium cacodylate pH overnight at 4°C. Cells were post-fixed with 1% osmium tetroxide and 1.5% potassium ferricyanide for 1 h, then with 1% uranyl acetate for 45 min, both in distilled water at RT in the dark. After washing, cells were dehydrated in graded ethanol solutions then finally infiltrated with epoxy resin and cured for 48 hs at 60°C. Sections of 70–80 nm thickness on formvar-coated grids were observed with a Hitachi H7500 TEM (Elexience, France), and images were acquired with a 1 Mpixel digital camera from AMT (Elexience, France).

### Scanning Electron microscopy (SEM)

Parasites were allowed to adhere and move on BSA coated-glass coverslips for 15 min at 37°C before being fixed with 2.5 % glutaraldehyde in 0.1 M sodium cacodylate for 30 min. After washing, cells were treated with 1 % osmium tetroxide in water, in the dark for 1 hour. Cells were dehydrated with increasing ethanol concentration baths. After two pure ethanol baths, cells were air-dried with HMDS. Finally, dry coverslips were mounted on stubs and coated with 5 nm platinum (Quorum Technologies Q150T, Milexia, France) and cells were imaged at 2 kV by a secondary electron detector with a Zeiss Merlin Compact VP SEM (Zeiss, France).

### Videomicroscopy

Time-lapse video microscopy was conducted in LabTek chambers installed on an Eclipse Ti inverted confocal microscope (Nikon France Instruments, Champigny sur Marne, France) with a temperature and CO_2_-controlled stage and chamber (Okolab), equipped with two Prime 95B Scientific Cameras (Photometrics, UK) and a CSU W1 spinning disk (Yokogawa, Roper Scientific, France). The microscope was piloted using MetaMorph software (Universal Imaging Corporation, Roper Scientific, France). A live-SR module (Gataca Systems, France) was added to the system to improve the obtained resolutions. Exposure time of 500 ms was used for the simultaneous acquisition of the GFP and mCherry channels, in dual camera mode (with band pass filters 525/50 nm and 578/105 nm, dichroic mirror at 560 nm, and laser excitation at 488 nm and 561 nm). Videos were captured at 2 frames per second (fps).

### Automatic Tracking and vesicle co-distribution using the Imaris Software

Automatic tracking of vesicles using the Imaris software (Bitplane, Oxford Instruments) was applied on the recorded videos retrieved from the GFP and mcherry channels of SAGΔGPI-GFP / mcherryRab11A-WT expressing parasites. We first used the tool “Spot detector” for selecting-filtering spot size and intensity values for each channel. Next, we manually removed detection of false GFP-positive spots (notably detected in the vacuolar space due to the secretion of the SAGΔGPI protein in Rab11A-WT parasites). The tool “Track Manager” was used to manually correct the obtained tracks when required and to extract the xy positions of a given spot over time enabling to calculate the Mean Square Displacement (MSD) using MATLAB (see below). The tool “spot co-localization” was used to calculate the percentage of co-distribution between DG and Rab11A-postive vesicles. A distance of 300 nm between the spots was selected corresponding to the average size of the vesicles. At a given time point and for the entire vacuole, the number of all detected green spots, as well as the number of green spots co-distributing with the red spots were extracted to calculate the co-distribution percentage. This was repeated over 5 consecutive time points every 2 s for the first 10 s of recording to avoid bleaching of the fluorescent signals. The mean co-distribution percentage over these 5 time points was calculated per vacuole. The mean +/− SD of 10 vacuoles was then calculated.

### Manual Tracking and Mathematical Modeling with MATLAB

When indicated, the manual tracking plugin from the Image J software (https://imagej-nih-gov/ij/) was applied on the images obtained with the MetaMorph software to extract in time the spatial xy positions of the fluorescent vesicles. In order to track and model the type of motion of the vesicle, images were processed in MATLAB (www.mathworks.com) by applying *fit* function (‘poly1’ or ‘poly2’ options).

MSD was calculated thanks to a MATLAB script according to the formula:

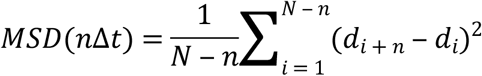

MSD curves were fitted according to the formula:

*MSD* = 4*Dt* + *v*^2^*t* (with D the Diffusion Coefficient and v the velocity), for directed motion

*MSD* = 4*Dt* (with D the Diffusion Coefficient), for normal diffusion.

### Statistics

Means and SEM / SD were calculated in GraphPad (Prism). *P*-values were calculated using the Student’s *t*-test assuming equal variance, unpaired samples and using two-tailed distribution.

## Acknowledgments

We thank Aoife Heaslip, Marc-Jan Gubbels, Dominique Soldati-Favre, Jean-François Dubremetz, Maryse Lebrun and Ali-Mohamed Hakimi for sharing of reagents. KV, SM and GL have been supported by the Laboratoire d’Excellence (LabEx) ParaFrap from the National Agency for Research ANR-11-LABX-0024 grant and by the ANR-14-CE14-0002-01 grant. SM has been supported by a joint Chaire d’Excellence from University of Lille and the Centre National pour la Recherche Scientifique (CNRS).

---

**Supplementary Figure 1**

**A-**Immunofluorescence assay showing the dense granule proteins GRA2 and GRA5 (green) retained in intra-cytosolic vesicles in Shield-1-induced (+S) Rab11A-DN expressing parasites, while being efficiently released into the vacuolar space and at the vacuole membrane in induced Rab11A-WT expressing parasites. The parasite cortex is delineated by GAP45 (red). Bars: 2μm. **B-** Fluorescence images showing the dense granule protein GRA16 (green) retained in intra-cytosolic vesicles in Shield-1-induced Rab11A-DN expressing parasites, while being secreted and translocated into the host cell nuclei (small arrows) in induced Rab11A-WT expressing parasites. Bars: 5μm. **C-** Immunofluorescence assay showing the localization of the non-secreted proteins GRA3 (red) and ROM4 (green) in distinct vesicles in Shield-1 (+ S) induced Rab11A-DN expressing parasites. In Rab11A-WT expressing parasites, both proteins are efficiently released at the vacuolar membrane and at the parasite plasma membrane. Bars: 2μm.

**Supplementary Figure 2**

**A**-Immunofluorescence assay showing the correct localization of SAG1, GAP45 and MLC1 in Shield-1-induced extracellular adherent Rab11A-DN expressing parasites. Bars: 2μm.

**Supplementary Figure 3**

Immunofluorescence assay showing the apical localization of MIC2-positive micronemes (A) and the plasma membrane protein ROM4 (B) in Shield-1 induced Rab11A-WT and Rab11A-DN parasites. Bars: 2μm.

**Supplementary Movie SM1 and SM2**

mcherryRab11A-positive vesicle (red) dynamics (left panel) in intracellular *T. gondii* parasites expressing IMC3-YFP (green). Imaging speed: 2 fps.

**Supplementary Movie SM3**

Movie showing mcherryRab11A-positive vesicles (left panel, arrows) moving along the developing daughter buds labeled with IMC3-YFP (right panel) during the process of cytokinesis. Imaging speed: 2 fps.

**Supplementary Movie SM4**

mcherryRab11A-positive vesicle (red) dynamics in intracellular *T. gondii* parasites expressing Cb-Emerald GFP (green) showing a mcherryRab11A-positive vesicles moving along cortical F-actin. Imaging speed: 2 fps.

**Supplementary Movie SM5**

Rab11A-positive vesicles (red) in close contact with dynamic cytosolic actin filaments in intracellular *T. gondii* parasites expressing Cb-Emerald GFP (green). Imaging speed: 2 fps.

**Supplementary Movie SM6**

mcherryRab11A-positive vesicle (red) dynamics in intracellular *T. gondii* parasites treated with cytochalasin D for 30 min before imaging. Imaging speed: 2 fps.

**Supplementary Movies SM7 and SM7b**

Movies showing the joint transport of a DG (green) docked on a Rab11A-positive vesicle (red) along the cortex of a SAG1△GPI-GFP and mcherryRab11A-WT expressing parasite. SM8b: tracking of the vesicles.

**Supplementary Movie SM8**

Automatic tracking of DG motion in SAG1△GPI-GFP expressing parasites.

**Supplementary Movie SM9**

Movie showing 3 DG tracks extracted from a region of interest of SM9 and analyzed for their mode of motion. Trajectory 2 (also shown in SM8) displays a directed motion, while trajectories 1 and 3 display confined diffusive motions.

**Supplementary Movie SM10**

Dense granule (green) dynamics in intracellular *T. gondii* parasites expressing SAG1 △GPI-GFP and mcherryRab11A-DN. The trajectories of 4 DG were tracked.

**Supplementary Movie SM11**

Dense granule (green) dynamics in intracellular *T. gondii* parasites expressing SAG1 △GPI-GFP and mcherryRab11A-DN 4h after Shield-1 removing in 0,5μM pre-induced Rab11ADN parasites. Imaging speed: 4 fps

**Supplementary Movie SM12**

Dense granule (green) dynamics in intracellular *T. gondii* parasites expressing SAG1 △GPI-GFP and mcherryRab11A-DN 4h after Shield-1 removing in 1μM pre-induced Rab11ADN parasites. Imaging speed: 2 fps.

**Supplementary Movie SM13**

mcherryRab11A-positive vesicle (red) dynamics in Shield-1 induced extracellular motile *T. gondii* parasite. Imaging speed: 2 fps.

**Supplementary Movie SM14**

mcherryRab11A-positive vesicle (left panel) dynamics in Shield-1 induced extracellular *T. gondii* parasite invading a host cell (right panel). Imaging speed: 2 fps.

